# Fast and Accurate Cell Tracking: a real-time cell segmentation and tracking algorithm to instantly export quantifiable cellular characteristics from large scale image data

**DOI:** 10.1101/2023.01.09.523224

**Authors:** Ting-Chun Chou, Li You, Cecile Beerens, Kate J. Feller, Miao-Ping Chien

## Abstract

Quantitative characterizations of cellular dynamics and features of individual cells from a large heterogenous population is essential to identify rare, disease-driving cells, which often exhibit aberrant cellular behaviors like abnormal division, aggressive migration or irregular phylogenetic cell lineages. A recent development in the combination of high-throughput screening microscopy with single cell profiling provides an unprecedented opportunity to decipher the underlying mechanisms of disease-driving phenotypes observed under a microscope. However, accurately and instantly processing large amounts of image data like longitudinal time lapse movies remains a technical challenge when an immediate analysis output (in minutes) of quantitative characterizations is required after data acquisition. Here we present a Fast and Accurate real-time Cell Tracking (FACT) algorithm, which combines GPU-based, ground truth-assisted trainable Weka segmentation and real-time Gaussian mixture model-based cell linking. FACT also implements an automatic cell track correction function to improve the tracking accuracy. With FACT, we can segment ∼20,000 cells in 2 seconds (∼4.5-27.5 times faster than state-of-the-art), and can export quantifiable features from the cell tracking results minutes after data acquisition (independent of the number of acquired image frames) with average 90-95% tracking precision. Such performance is not feasible with state-of-the-art cell tracking algorithms. We applied FACT to real-time identify directionally migrating glioblastoma cells with 96% precision and to identify rare, irregular cell lineages in a population of ∼10,000 cells from a 24hr-time lapse movie with an average 91% F1 score, results from both were exported instantly, mere minutes after image acquisition.

## Introduction

The advent of modern, large-scale volumetric fluorescence microscopy and high-throughput screening microscopy have driven rapid development in computational methods^1,2^. Several advanced, automated computational methods have been recently introduced to accurately and rapidly process the big multi-dimensional image data generated from these new microscopes^1-4^ and to extract meaningfully biological information: cell migration, division, morphological changes, differentiation or cell-cell interactions^1-3,5,6^.

Prior to these newly developed computational methods, terabytes of image data require months of processing instead of a number of days^7-9^. Real-time Accurate Cell-shape Extractor (RACE)^1^, for example, can segment cells from 3,836 time points within 1.4 days compared to 100-700 days used by other state-of-the art cell segmentation methods^7-9^. To accurately and rapidly process cellular dynamics and lineages, Tracking Gaussian Mixture Model (TGMM) cell tracking algorithm^2^, for another example, can process 26,000 cells/min (including cell segmentation + cell linking), compared to other state-of-the art cell tracking algorithms, which can only process a number of thousands of cells/min^10,11^.

Inspired by TGMM, our recently developed cell tracking algorithm, mTGMM (modified TGMM)^3^, can process ∼42,000 cells/min and which has been successfully applied to process and extract quantifiable characteristics and phenotypic information of individual cells (i.e., cells with fast migration or spindle-shaped morphology)^3^ imaged through our high-throughput screening microscope, Ultrawide Field-of-view Optical (UFO) microscope^3^. mTGMM can efficiently process UFO-curated big image data, for instance, 20,000 cells/frame of a 2hr-time lapse movie (less than 30 frames) within 10 minutes. This processing time is sufficient for subsequent, selective isolation of cells of interest from the imaged dish (for cells migrate ≤ 1 μm/min) as those target cells remain closely to the coordinates of the last acquired image frame. The capability to rapidly process and isolate subpopulations of cells from a large pool of heterogenous cells opens an unprecedented possibility to dissect molecular mechanisms of phenotypes of interest, as the isolated cells with desired phenotypes can be directly subjected to downstream (single cell) sequencing and profiling. The profiled genomes, transcriptomes or proteomes can be straightforwardly linked to their corresponding cellular phenotypes (namely, genotype-to-phenotype linking)^3,12,13^.

Despite the remarkable advances in fast cell tracking methods (cell segmentation + cell linking), these methods decrease tracking accuracy when cells have complicated cellular behaviors (like cells migrate across each other) or cells are densely populated (> ∼100,000 cells/cm^2^); such a density often occurs when tracking cells or studying cell lineages over days or is required when searching for rare biological events (0-5 %). Most importantly, the processing speed of these new computational methods cannot satisfy the demand of instantly processing hundreds or thousands of image frames within a number of minutes, which is critical when isolating cells of interest from the imaged dish is required. These constraints stop us from directly profiling cells displaying among others abnormal, longitudinal migratory behaviors, anomalous cell lineages (both require longitudinal imaging) or irregular cell divisions (rare biological events), which are important in cancer research and treatment development. Deciphering the underlying mechanisms of these abnormal phenotypes is important in this context as the occurrence of cells exhibiting these characteristics usually linearly correlates with poor prognosis^14-16^.

To have superior cell linking and tracking performance, high accuracy in cell segmentation is extremely critical. Deep neuronal network-based cell segmentation have been demonstrated to address this challenge^17^, but the amount of labeled data required for training to reach high performance (> 90% of accuracy at intersection-over-union (IoU) threshold of 0.5) often limits the widespread applications of these methods; this problem is intensified when applying to different types of cells or samples as new training is often required.

To address the abovementioned challenges, here we present FACT (Fast and Accurate real-time Cell Tracking algorithm). FACT is a real-time, instant cell segmentation and cell linking algorithm combining GPU-based, machine learning-assisted cell segmentation (Figure 1) and Gaussian mixture model-based cell tracking algorithm with automatic cell track correction to accurately and instantly process big image data (i.e., 20,000-30,000 cells/frame for > hundreds of image frames) and export quantitative characteristics of individual cells minutes after data acquisition. The algorithm can easily be adapted to segment different cell types with 30-60 minutes of human annotation to achieve peak performance for unseen sample types and have higher segmentation and tracking performance than state-of-the art.

**Figure 1.**
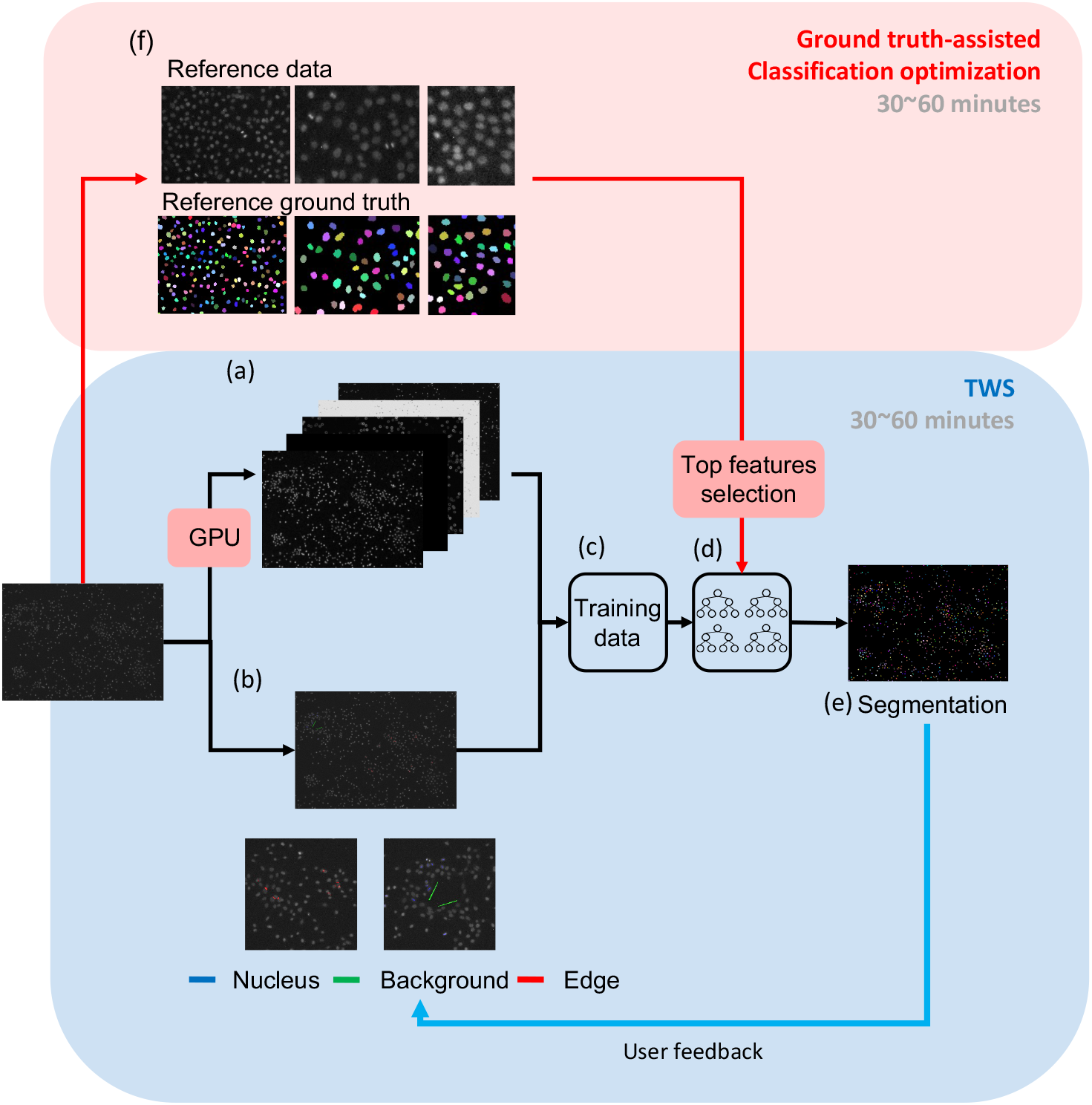
Ground truth-assisted Trainable Weka Segmentation (GTWS) pipeline. The original TWS pipeline is highlighted in blue: (a) Image features, generated by applying all default image filters on an input image. (b) Data annotation (or labeling). Manual assignment of pixels to three classes: nucleus (blue), background (green) and edge(red). (c) Training data, a combination of annotated pixels and their corresponding values from image features. (d) Random forest classifier, which is trained with the input training data. (e) We then use the well-trained model to perform the semantic segmentation. Connected component analysis is applied to generate the instance segmentation from the semantic segmentation. If needed, users can improve the segmentation performance by adding more annotations and re-training the classifier (Step a-d) until no further improvement is observed. The processes from (a) to (e) take ∼ 30-60 minutes. Our GTWS (highlighted in red) improves the segmentation speed via GPU-based operation and key feature selection and increases the segmentation accuracy by incorporating ground truth reference data (f), which only requires 30-60 minutes of preparation. The reference data is used to optimize the random forest classifier. Reference data are generated from a number of cropped regions of the original input image and labelled as individual nuclei and background.

### Cell segmentation with sparse annotation and high accuracy with FACT

As mentioned, deep learning-based models^17^ (i.e., U-NET^18^ or mask R-CNN^19^) have been shown to perform well when used to tackle difficult problems in cellular image analysis and image segmentation^17^. However, creating or (re-)training models that are customized for each sample type or experiment is often required to reach high performance, resulting in hours to days of data annotation, ground truth preparation and model training. Low annotation machine learning methods, such as Trainable Weka Segmentation (TWS)^20^ or ilastik^21^, on the other hand, can finish data annotation and training for each sample type within 30-60 minutes, and therefore are easier to implement when model creation or (re)training is required. Despite the great potential and promising results of TWS or ilastik, a better cell segmentation performance (i.e., >95% F1 score at IoU threshold of 0.5) is required to have higher accuracy in cell tracking and cell lineage construction. The necessary performance increases seem difficult to reach using TWS, ilastik or even deep learning methods^17^. In addition, TWS and ilastik require processing time of ∼55 seconds or ∼20 seconds per image frame, respectively (for a 4096 × 3000 pixels image, with ∼10,000-30,000 cells/image; Figure S1). A shorter processing time (less than 5 sec/frame) is often required when screening and identifying rare biological events from a large quantity of cells, especially when immediate isolation of those rare cells is needed.

To increase segmentation performance and to improve processing time per frame, here we introduce a new GPU-based, ground truth-assisted TWS method, called Ground truth-assisted Trainable Weka Segmentation (GTWS) (Figure 1). GTWS improvs cell segmentation accuracy (> 95% F1 score at IoU threshold of 0.5, Figure 2 and S2) by optimizing random forest classification using sparse ground truth reference data, which consists of foreground and background labelled images and which can be prepared within 30-60 minutes (Figure 1, Figure S3, S4 and **Materials and Methods**). The implementation of the ground truth data acts as a reference during the process of selecting the best random forest parameter option (i.e., the depth of the tree and the number of leaves etc.) by checking the segmentation performance (i.e., accuracy or F1 score) (Figure 1, see Figure S3 for detail).

**Figure 2.**
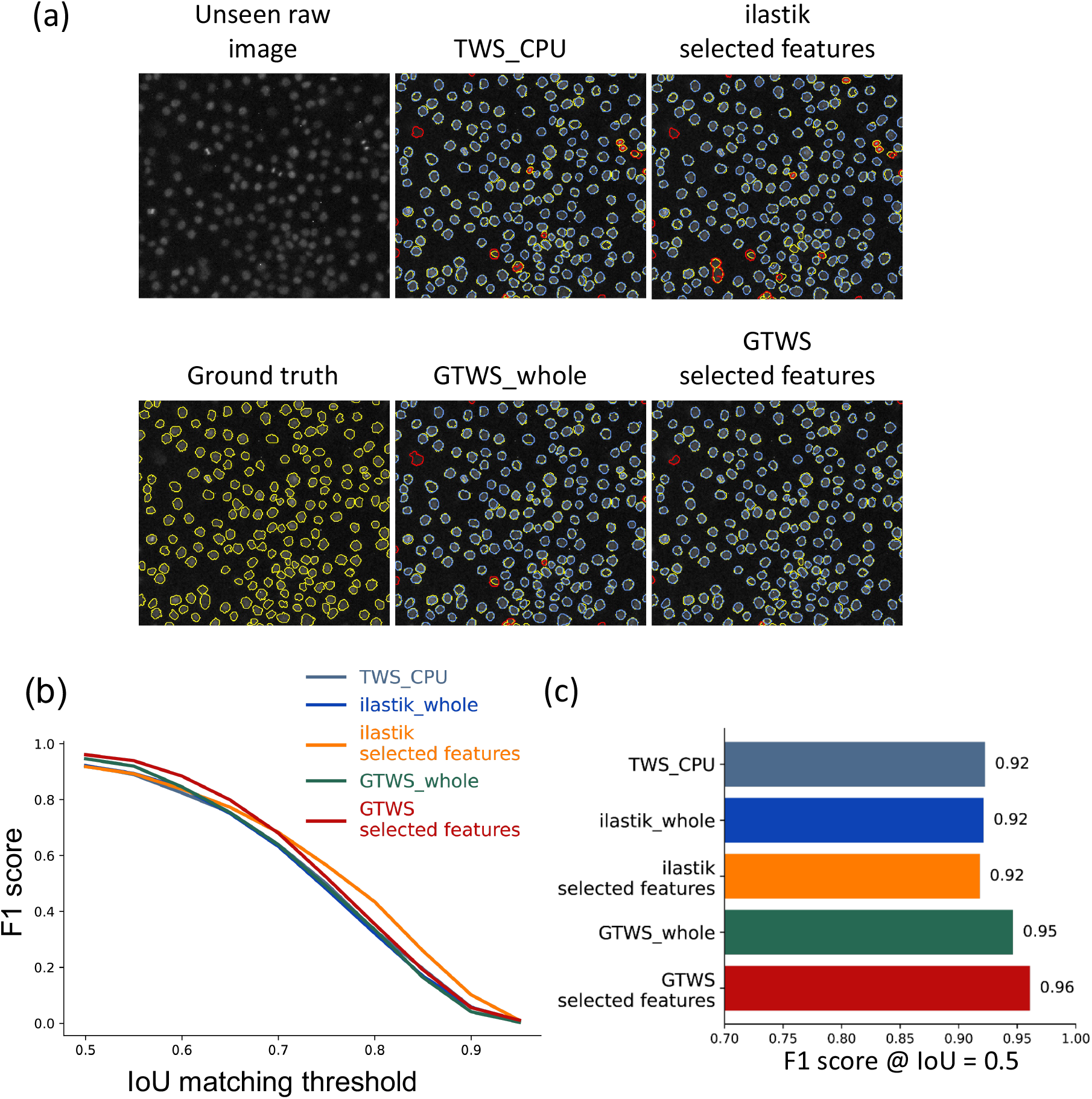
Benchmarking of Weka-based cell segmentation methods (TWS, ilastik and GTWS). Cell segmentation from different Weka-based segmentation methods: TWS (top middle), ilastik’s key feature selection method (via the “filter” feature selection method (3 features)) (top right), GTWS’ whole image features (GTWS_whole; bottom middle) and GTWS’ top selected features (3 features) (bottom right). These outcomes were achieved using the same input image (top left). Ground truth (GT) segmentation (bottom left) was prepared independently (manual segmentation). Object contours are shown in yellow. Each segmentation outcome (per method) was compared to the GT. In each of the 4 segmented images, we color-coded the results: cells in blue indicate a satisfactory outcome (when calculated IoU ≥ 0.5); cells in red are false-positive (when calculated 0.1≤ IoU < 0.5); cells in yellow are false-negative (when calculated IoU < 0.1). (b) F1 score of each method when using different IoU values as a threshold. The curve is obtained as follows: each IoU threshold ∈ {0.5, …, 0.9} is considered as a threshold to calculate segmented true-positive (TP), false-positive (FP) and false-negative (FN) values; A cell’s segmentation is compared to its GT, and it is a true-positive if the calculated IoU ≥ IoU threshold; a false-positive if the calculated IoU, 0.1 < IoU < IoU threshold; a false-negative if the calculated IoU < 0.1. Given all TPs, FPs and FNs values, we calculated the F1 score, shown as the y-axis. (c) Averaged F1 score of each segmentation method when using the IoU threshold of 0.5.

The minimal endeavor of ground truth preparation makes the method easily adaptable to other sample types without compromising its performance. Furthermore, GTWS improves the processing time by i) computing the segmentation via GPU instead of CPU (**Materials and Methods**, Figure 1) and by ii) selecting the most relevant image features for classification (instead of running all 57 image features)^20,21^ (Figure 2 and S2). Image features are the images processed with a number of different filters (Gaussian filer, Sobel filter, difference of Gaussians filter, Hessian matrix, Membrane projections and Median filter). They hold major information per image, e.g., a Sobel filter can find out pixels that represent a nuclei’s edge, while suppressing the edge-irrelevant pixels. These image features will then feed into a random forest model. The ground truth reference data, used in the selection of the best random forest model, is also applied to the selection of the most relevant image features (See Figure S3 for detail). By default, TWS generates 57 image features (whole features) and uses them to construct a random forest model. With our GTWS, we observed that a model with a subset of image features (selected features) yields faster and more accurate prediction (Figure 2 and S2), due to the fact that not all features are relevant to the data (Figure S5). We implemented forward selection as our key feature selection method^22^ (see **Materials and Methods** for detail) and found that a model with 3 key selected features yields as similar performance (96 % F1 score) as whole features (95 % F1 score) (Figure 2), but computes much faster (< 2 sec/frame) than those using larger than 3 features (Figure S6). With this processing speed, we can, for example, identify extremely rare tripolar dividing cells: ∼50 division events, resulting in 3 cells per division for a total of ∼150 cells, from a population of ∼0.8 million MCF10A breast cancer cells (∼0.05-0.1% occurrence rate) in ∼5 minutes after data acquisition (∼100 seconds of cell segmentation + ∼200 seconds of tripolar division detection) (Figure S7). Identifying larger than ∼100 cells of interest (from a cell population) is essential for experiments involving downstream single cell isolation and sequencing^3,13^, and profiling > ∼100 cells is required to draw a statistical conclusion: this makes it necessary to screen, segment and identify individual cells in a population of close to a million cells in a matter of minutes which GTWS makes possible.

ilastik^21^, arguably the most commonly used, state-of-the art Weka-based cell segmentation method^20^, also allows users to select key features, instead of using its whole features (32 image features are used in ilastik, **Materials and Methods**). ilastik offers three different types of key feature selection methods, namely filter method, GINI importance and wrapper method (Figure S2), none of these feature selection methods outperforms either our GTWS feature selection method nor ilastik’s whole feature method (Figure S2).

By optimizing random forest classification using sparse, reference ground truth data and improving the processing time via GPU and key feature selection, our GTWS outperforms state-of-the art TWS and ilastik methods, both in segmentation performance (96% vs 92% F1 score) and computational time (27.5 times faster than TWS and 4.5-10 times faster than ilastik) (**Materials and Methods**, Figure 2, Figure S1 & S2). Please note that the segmentation performance was evaluated based on the comparison of ground truth data (of unseen images) and the segmented result (Figure 2). The high performance on unseen data indicates that our model was not overfitted.

### Instant cell linking and cell track correction with FACT

A highly accurate cell segmentation is required and crucial for accurate cell tracking. With the high performance of our GTWS cell segmentation, we then apply it to link and track cells from frame-to-frame. The ability to accurately link positions of individual cells between frame-to-frame and to reconstruct the movement and divisions of cells is crucial for dissecting cell lineage and their correlation with cell functions and cell-fate decisions^16,23^. TGMM^2^ and mTGMM^3^ are fast state-of-the art cell linking and tracking algorithms as ∼26,000 and ∼42,000 cells can be processed per minute, respectively, compared to other cell tracking algorithms, which can only process a number of thousands of cells/min^10,11^. Tracking cell lineages often requires over a day of image acquisition, resulting in hundreds to thousands of image frames. Even with mTGMM, the algorithm cannot sufficiently process this amount of image frames within a number of minutes, which is required for downstream phenotype-to-genotype linking experiments. We therefore develop an instant, Gaussian mixture model-based cell linking algorithm (Figure 3). Similar to TGMM^2^ and mTGMM^3^, individual cells are modelled as a 2D Gaussian, 𝒩(*x*; *μ, Σ*) with *x* be the 2D coordinates, *μ* be the mean location and Σ be the covariance (i.e., representing the shape of a Gaussian blob). All Gaussians (representing all individual cells) of the entire image constitute a Gaussian mixture per image frame. Calculating the expected properties per cell (i.e., intensity, size, shape) over time is performed by a full Bayesian approach^2,24^, with the assumption that cell properties between two consecutive frames are correlated. If correlated, the same cells of the current frame and the previous frame will be linked (see detail in **Materials and Methods**). To instantly complete image analysis (independent of the number of frames) after the acquisition of the last image frame, we implemented a real-time cell segmentation and linking method (Figure 3): once a new image is generated, it is directly processed by our GTWS cell segmentation; when two consecutive images are segmented, cells of these two image frames are linked and tracked; lastly, when the tracked frames are > 3, the algorithm reviews all the connected tracks (from the current frame and a past few frames, pre-defined by a window size, i.e., a number of frames) and perform cell track corrections if needed (see below). With this instant processing of freshly generated image frames, FACT can complete the cell segmentation and linking and export quantifiable features in a matter of minutes, independent of the number of image frames. In our case, processing a 24 hr-time lapse movie (361 frames with 4 minutes interval, ∼10,000 cells per frame) took 2.88 hours after data acquisition to export quantifiable results by mTGMM, while FACT required 0.17 hours to obtain the results. FACT is built based on mTGMM^3^ with further development of real-time cell linking and cell track correction. FACT’s computational speed maintains ∼30,000 cells/min, which tends to be slower than mTGMM as we add the function of cell track correction. Accurate cell lineage reconstruction highly relies on the precise detection of cell division and the differentiation of true divisions from false divisions. False cell divisions often arise from either mis-segmented cells or cells crossing each other’s trajectory; methods to correct the former cases have been reported^23,25,26^, but not for the latter scenarios. When two cells move into each other and share a fraction of their boundaries, this causes the disappearance of one cell track (Figure 4). We call this phenomenon cell merging. A false cell division occurs when the two cells demerge, meaning the two cells move apart from each other after merging (Figure 4). Figure 4b shows cell merging-demerging found in the glioblastoma (GBM) image data. The cell merging-demerging problem is exacerbated when cell density is high or cell behavior is complicated such as directionally walking pattern employed by GBM cells (Figure 4b). To address this challenge, FACT implemented an automatic, real-time cell track correction function, which includes three steps as follows: i) When a cell division is detected, we search for any incomplete cell tracks nearby (Figure 5a, top). Should the division occur next to an incomplete cell track within a pre-defined (Euclidean) distance (i.e., 10 pixels) and time window (i.e., 5 image frames), the division is deemed false division. The pre-defined distance and time window are based on user’s prior knowledge for each cell type. ii) When a false division is detected, the link between the mother and daughter cells will be disconnected and the falsely generated cell link will be removed accordingly (Figure 5a, middle). iii) Two incomplete tracks (after Step ii) will be re-connected by generating “fake” cell(s) in any position between the two incomplete tracks (Figure 5a, bottom).

**Figure 3.**
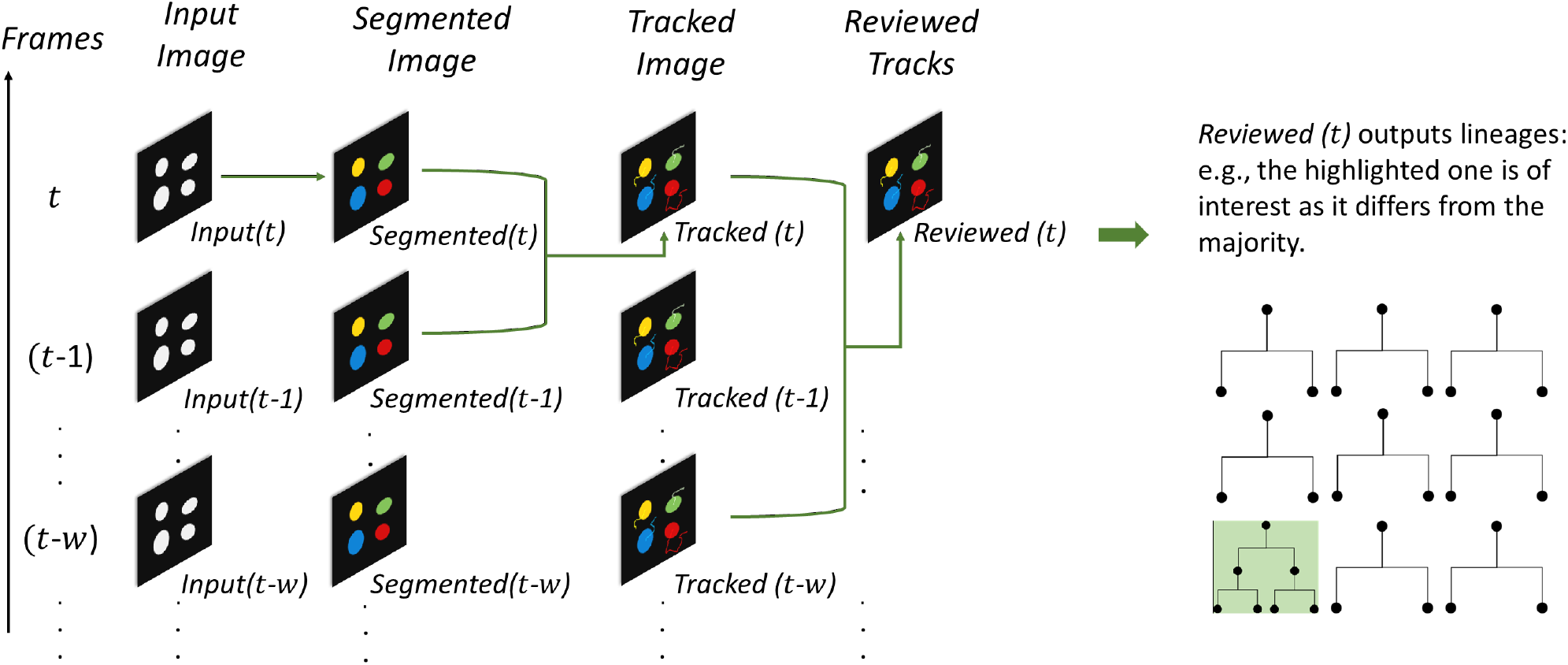
An overview of FACT pipeline to instantly segment cells and track cell lineages. Let’s assume in total *T* frames are required. At every frame-up-to-date *t* ∈ {1,2, …, *T*} a microscopy image *Input*(*t*) is generated (top row). It is immediately followed by GTWS segmentation, generating a segmented image *Segmented*(*t*). This segmented image *Segmented*(*t*) is linked to *Segmented*(*t* − 1) via tracking a short time lapse of two adjacent frames (*t* − 1) and *t*. Tracking is also up-to-date at *t*, i.e., *Tracked*(*t*), but likely with errors when some cells are mis-linked over time. A review step applies right after to examine all tracked cells from frames (*t* − *w*) to *t*, with *w* be a frame window size. The review particularly looks for the following errors: 1) incomplete tracks, i.e., cells that are not segmented/tracked at every frame, and 2) divisions caused by cell merging and demerging (see Figure 4-5). These errors could be corrected, to a certain degree, through a review over (*w k* 1) frames. We use *Reviewed*(*t*) to denote all tracks that are reviewed up to frame *t*. When *t* = *T* we reach the tracking result *Reviewed*(*t*) of a long-term imaging, with *Reviewed*(*t*) gives also the lineage information over time. Lineage that is different from the majority is captured as the abnormal one (i.e., the highlighted lineage tree has more divisions than the rest), representing our lineage of interest for downstream analysis. The review step could be disabled by setting *w* = 0, then *Tracked*(*t*) would represent the tracking result.

We verified our cell track correction method on GBM imaging data (Figure 5b, c), as these cells often travel directionally from point to point^27^, and the cell merging-demerging issue is exacerbated in this cell type. For example, when tracking ∼3000 cells over 5 hours, ∼450 merging-demerging events were detected. With our cell track correction method, we have corrected those (∼450) falsely detected cell division: with correction we reached 91% precision, while without correction the precision is 12% due to heavy contamination from merging-demerging.

**Figure 4.**
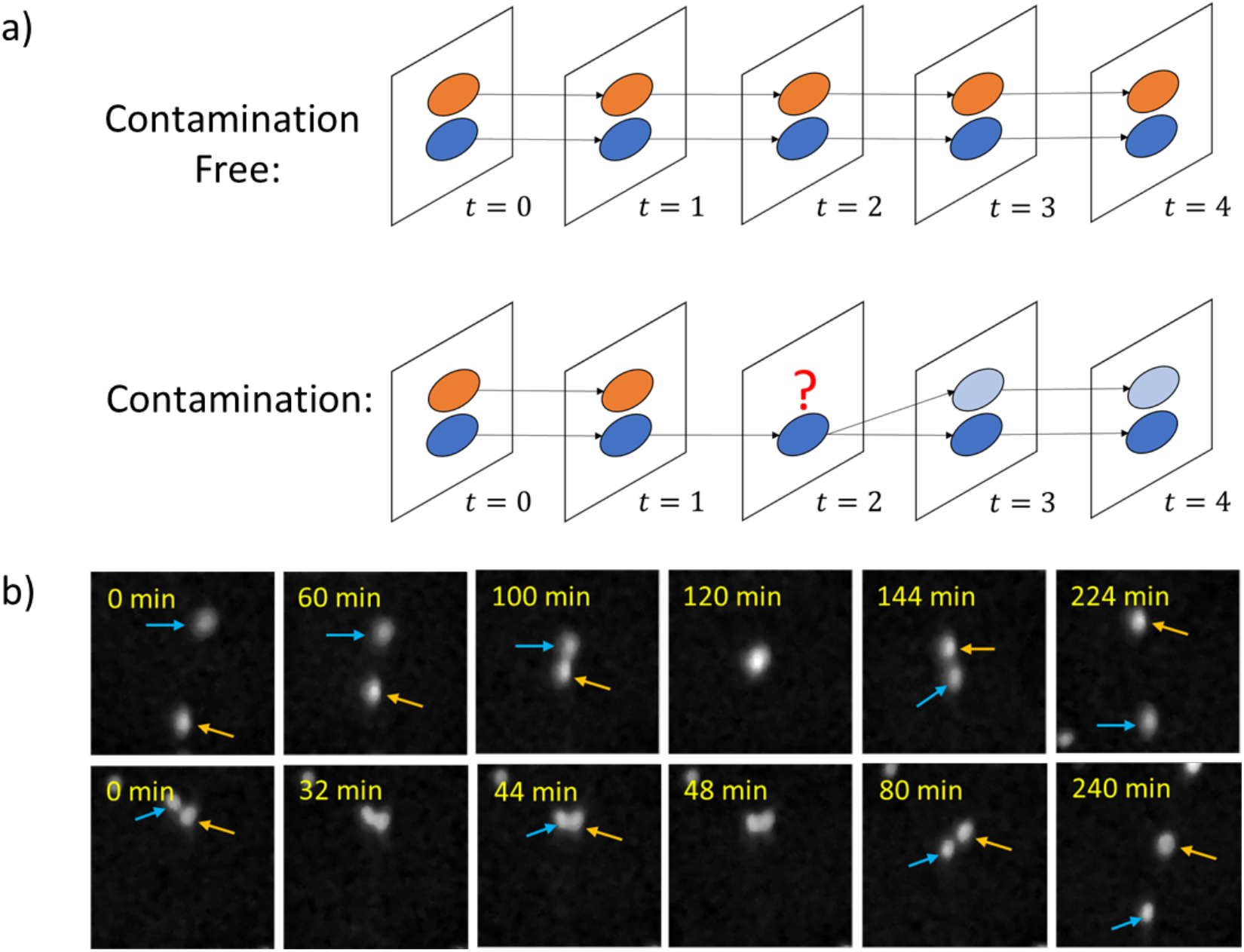
Cell merging and demerging. a) Graphical illustration of two complete tracks (top) and incomplete tracks (bottom). For the contaminated case, two cells merge at frame *t* = 2, giving rise to i) a false division at frame *t* = 3, and ii) an incomplete track of the orange cell that ends at frame *t* = 1. b) Two examples of the merging-demerging problem shown in the raw image data of glioblastoma (GBM) cells. Top: two cells (blue and orange arrows) merge for over 30 minutes, and deviate. A snapshot at time “120 min” shows the merging event. Bottom: two cells (blue and orange arrows) merge twice at time “32 min” and “48 min”. This is caused by both complicated cell behaviors as well as false segmentation. Time lapse videos can be found in Supplementary Video 1 and 2.

**Figure 5.**
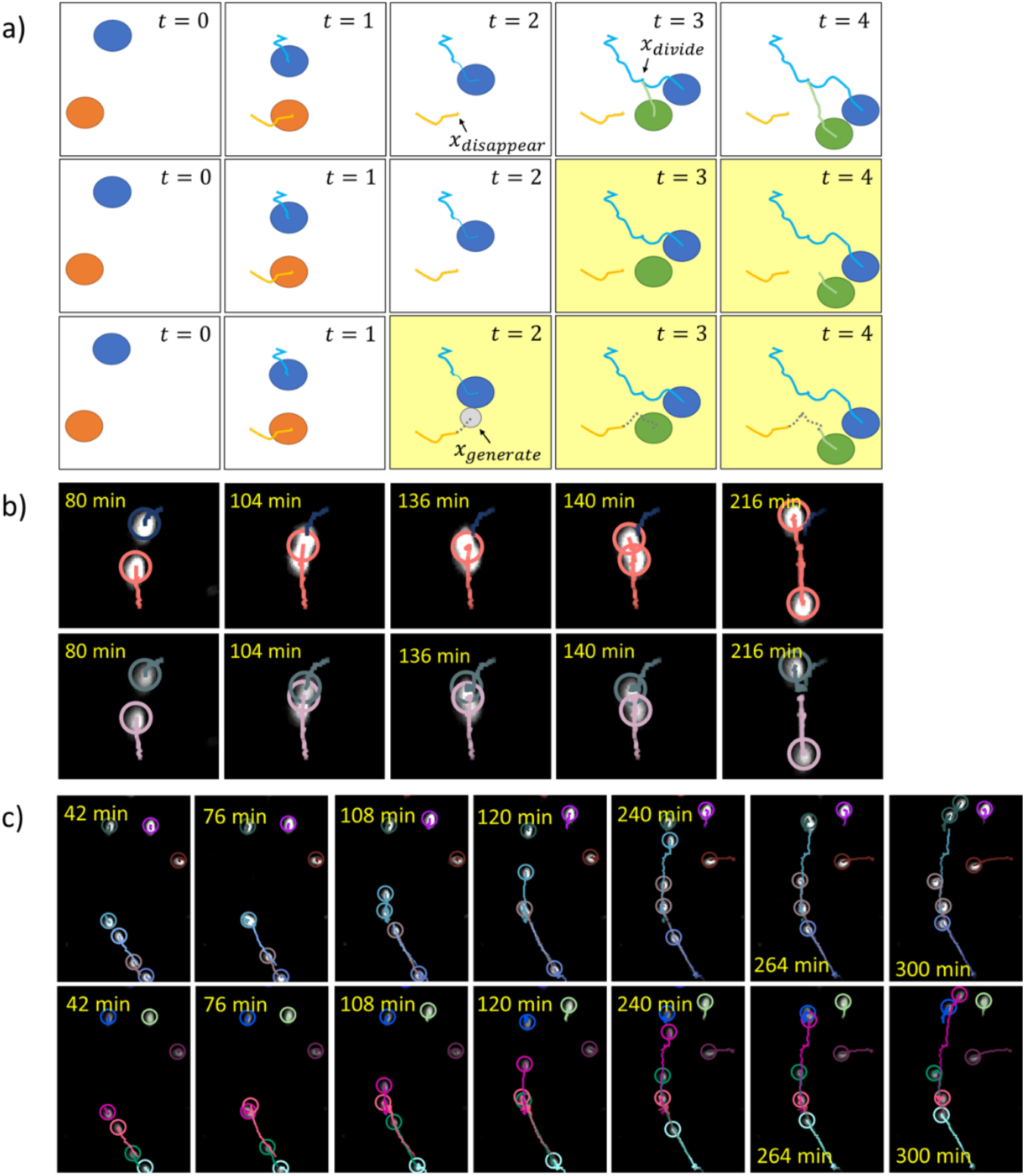
Cell track correction. a) Graphic illustration of cell track correction. **a)-top, Step i)** – to examine if a division is valid. We see that a merging at *t* = 2 causes a cell track (orange) disappearing at location *x*_*disappear*_, a demerging at *t* = 3 causes a division at location *x*_*divide*_. The false cell division is detected when the event is next to an incomplete track within a pre-defined (Euclidean) distance (Δ*x* between *x*_*disappear*_ and *x*_*divide*_) and within a pre-defined time window (time difference, Δ*t*, between the merging and demerging events). **a)-middle, Step ii)** – to remove the link of a false division. Compared to Step i), the frames with changes are highlighted in yellow. We disconnect the link (green) from *t* = 2 to *t* = 3 (in Step i), as this daughter cell (green) is closer to *x*_*disappear*_ than the other daughter cell (blue). The link removal generates an incomplete track (green) from *t* = 3 to *t* = 4. **a)-bottom, Step iii)** – to close the gap between the incomplete tracks. Compared to Step 2, the frames with changes are highlighted in yellow. The two incomplete tracks (orange and green) are to be connected. There is a gap between them, which refers to the disappeared cell at *t* = 2. We then construct a fake cell (gray) at this frame at any location of *x*_*generate*_ between *x*_*disappear*_ and *x*_*divide*_. We update the reconstructed cell to the subsequent frames *t* = 3, 4. At *t* = 4 we obtain two complete tracks. b) Merging-demerging caused a false division (top). The tracks are corrected with our cell track correction method (bottom). c) Merging-demerging caused three false divisions (top) (merging happened at ‘76 min’, ‘120 min’ and ‘264 min’), and tracks were corrected with our cell track correction method (bottom). Tracked and corrected videos can be found in Supplementary Videos 3 and 4.

### FACT identifies abnormal cancer cells

We then applied FACT to identify abnormally migrating GBM cells, namely the most directionally migrating cells as this behavior is linear correlated with cancer cell aggressiveness^28-30^. We used the metric of directionality ratio 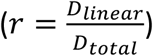 to measure the deviation between the cumulative distance *D*_*total*_ (total distance travelled over all time points) and the linear distance *D*_*linear*_ (the distance from the start to the end time point). When *r* is close to 1, a cell’s walk is close to directional walk. From a 5 hr-time lapse movie (see the time lapse movie in Supplementary Video 5), ∼3000 GBM cells were tracked and analyzed via FACT. Please note that manual checking and validation of all the tracked trajectories are required, and therefore a relatively small number of cells were used in this particular application. The directionality ratio (*r*) of all the tracked cells was processed immediately and exported in <2 minutes after data acquisition. Based on the density distribution of all directionality ratios (Figure 6), the mean ratio *μ* (0.40) and the standard deviation *σ* (0.24) were calculated. We set the ratio *r* > 0.90 as a threshold to define a directional trajectory. In this case we found 28 single cells (∼1.0% of the whole population) displaying directional walking pattern after the FACT analysis and we have validated the identified result manually with a precision of 96% (Figure 6 and Figure S8).

**Figure 6.**
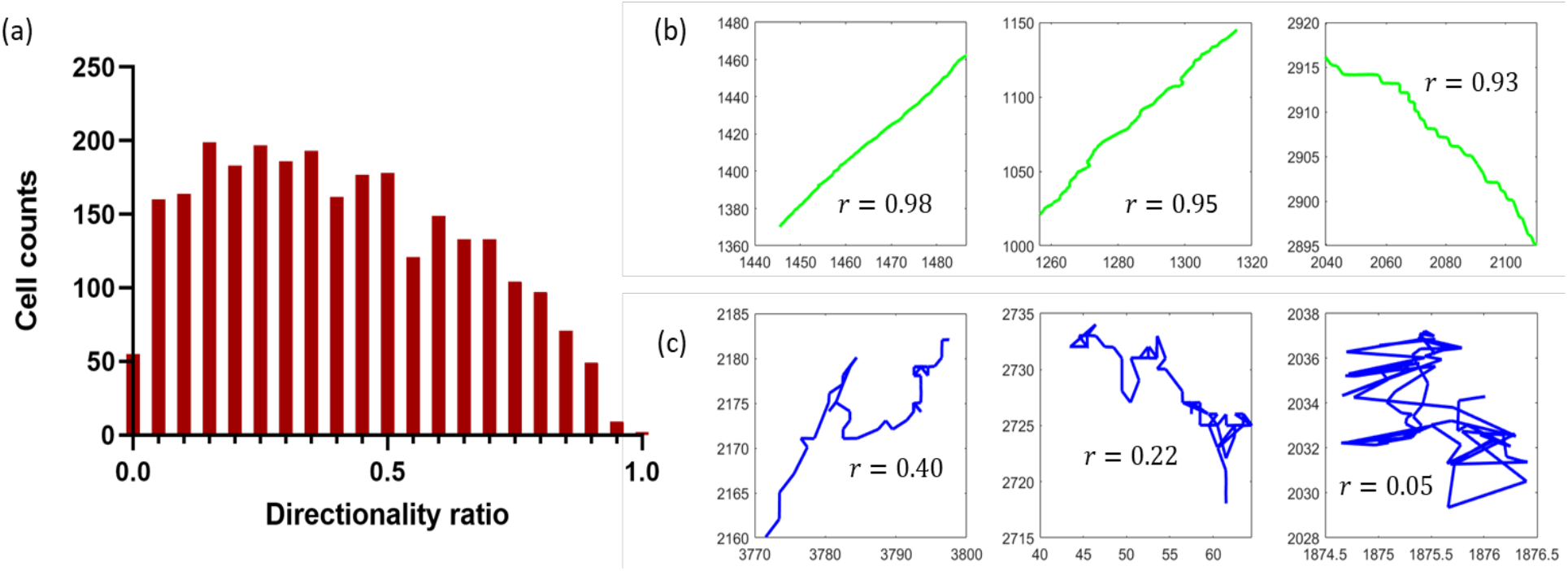
Application of FACT to GBM cell tracking. (a) Distribution of directionality ratio (*r*) over 2724 cells. Over all cells we looked for the ones giving ratio *r* > 0.90. (b) Example trajectories of cells with directional walk, directionality ratio (*r*) is indicated per cell. (c) Example trajectories of cells without directional walk, directionality ratio (*r*) is indicated per cell.

Next, we applied FACT to real-time segment and track ∼10,000 MCF10A cells from a 24 hr-time lapse movie (4 min/frame and 361 frames in total, Supplementary Video 6) (Figure 7 and Table S1), from which cell lineages of individual cells will be extracted. We aim to identify abnormal cell lineages which deviate from the rest of lineages. MCF10A cells are epithelial breast cancer cells with tendency to densely pack and connect with each other, and are therefore used as the sample model here.

**Figure 7.**
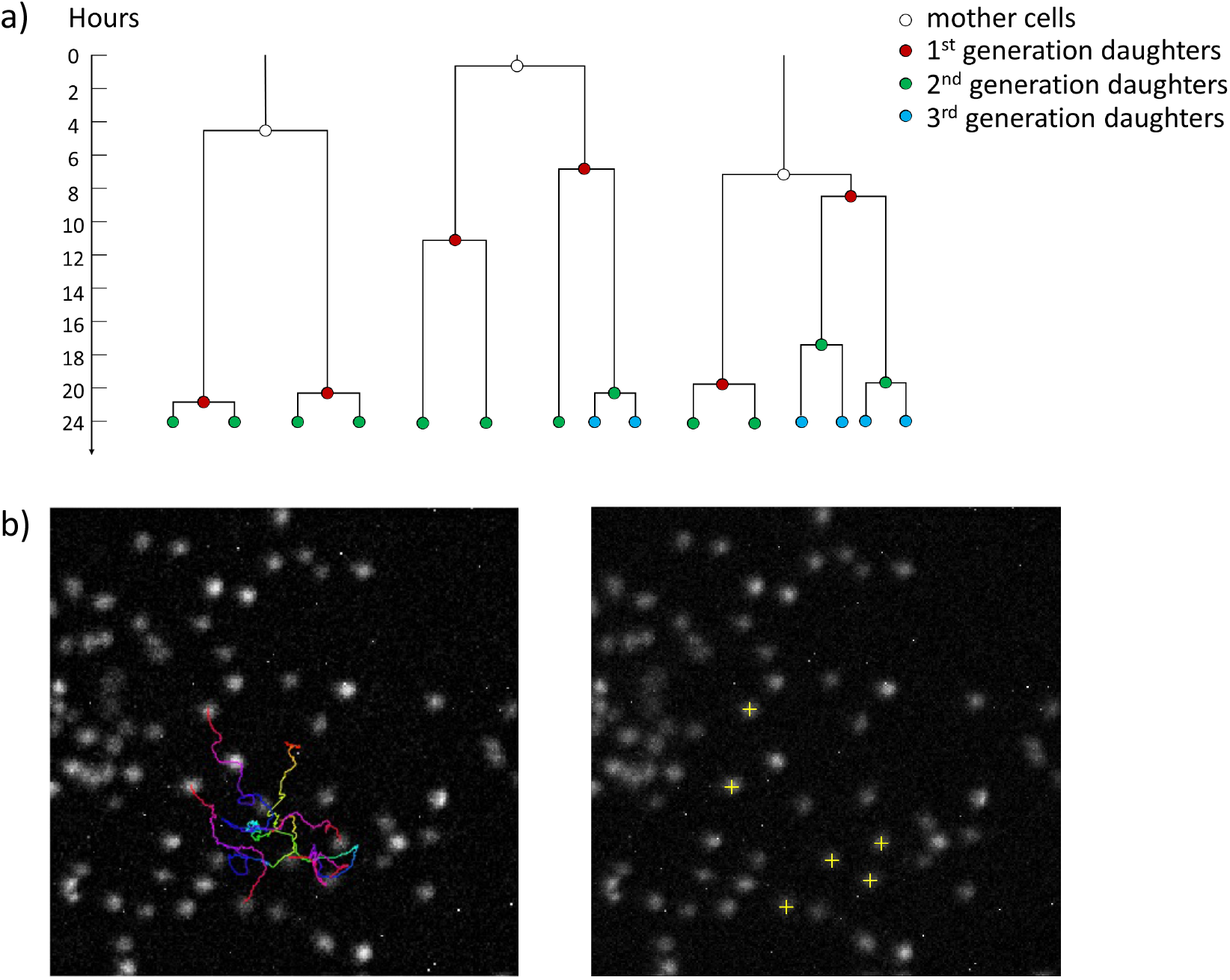
Cell lineage tracking of MCF10A cells. a) Topology of lineages from 3 groups, ‘tree-3-div’ (left), ‘tree-4-div’ (middle) and ‘tree-5-div’ (right). In each plot, time is represented as the vertical axis, going from 0 to 24 hours. Daughter cells generated in different generations are color-coded. Videos of these cases are included in Supplementary Video 9. b) Migratory trajectories of the lineage of interest (tree-5-div) (left image), and the final coordinates of the cells (yellow cross, right image).

After FACT, we grouped the trees by the number of cell divisions, namely, ‘tree-1-div’, ‘tree-2-div’ to ‘tree-5-div’, indicating a tree of 1, 2 to 5 cell divisions, respectively (Figure 7). More specifically, cells in the ‘tree-3-div’ group (as an example) contain 3 divisions within 24 hours. From the assayed ∼10,000 MCF10A cells, FACT exported the lineage information of each cell less than ∼10 minutes after the data acquisition with an average of 91% F1 score. We identified 52.8%, 12.2%, 6.9%, 0.5% and 0.1% of cells dividing once, twice, three, four and five times, respectively. Figure 7a shows 3 representative lineage trees of fast dividing cells (‘tree-3 to 5-div’) identified from the assayed cells (see the original movies of these 3 lineages in Supplementary Video 7-9), from which the ‘tree-5-div’ case is shown in Figure 7b. The coordinates of those 5 daughter cells of this irregular lineage tree were immediately exported within 10 minutes after the image acquisition.

In conclusion, we have developed a real-time cell segmentation and tracking algorithm, FACT, to instantly process large scale image data (> 10^4-6^ cells/frame for > 10^2-3^ image frames) and export quantitative cellular characteristics within minutes after data acquisition. FACT implements ground truth-assisted Weka based cell segmentation (GTWS), which requires only 30-60 minutes of human annotation to achieve peak performance for unseen sample types. High segmentation performance (>95% F1 score at IoU threshold of 0.5), in combination with real-time cell track correction, results in high tracking performance (90-95% precision) as the program can correct wrongly detected cell divisions and falsely linked cell tracks in a real-time fashion. As shown, FACT can correctly track GBM cells, which tend to migrate across each other, and track densely packed MCF10A breast cancer cells, with on average 90-95% precision.

When combined with high-throughput screening microscopy, FACT can be used to instantly identify rare subpopulations of cells during large scale image data acquisition, enabling immediate isolation of target cells for downstream assays, like single cell sequencing or proteomic profiling^3,13^. For example, FACT can instantly identify sparse, tripolar dividing cells, aggressively migrating GBM cells, and fast dividing cells, all extremely low occurrence rate events. Linking rare abnormal cellular phenotypes (i.e., metastatic cells or abnormally dividing cells) to genotypes, transcriptomes or proteomics is possible with FACT as instant export of cellular characteristics is required, to enable target cell isolation; this ability creates an unprecedented opportunity to decipher the underlying mechanisms of the observed irregular, disease-driving phenotypes.

## Materials and Methods

### Imaging data preparation

The images used in this paper are either MCF10A breast cancer cells or glioblastoma (GBM) primary culture cells (see Supplementary Methods for the detail of cell culture). Images were taken by a custom-built microscope, ultrawide field-of-view optical (UFO) microscope^3^, which incorporates a large chip-size CMOS Point Grey camera (GS3-U3-123S6M-C, 4,096 × 3,000 pixels, 3.45 μm per pixel, FLIR) and large-field-of-view (FOV) objectives (Olympus MVP Plan Apochromat, ×0.63)^3^ with comparatively high numerical aperture (NA = 0.25).

### Workstation and software information

All segmentation work was done via Ubuntu 18.04 environment with Intel(R) i9-9980XE CPU, 128 GB ram, two of Nvidia QUADRO RTX 8000 GPUs. GTWS was implemented in Python 3 using open-source packages: NumPy^31^, OpenCV^32^, CuPy^33^, cuML^34^, scikit-image^35^, SciPy^36^ and Matplotlib^37^.

Real-time cell linking was performed on a single workstation, Dell Precision 7920, with the following hardware components: dual Intel(R) Xeon(R) Gold 6130 CPUs, 12×32GB ram, a Nvidia GeForce GTX 1080 GPU. The FACT software will be accessed in GitHub after paper acceptance.

### Details of Ground truth-assisted Trainable Weka Segmentation (GTWS)

GTWS is inspired by and modified from original *Trainable Weka Segmentation* (TWS)^20^. We first introduce the method, and we introduce and explain our improvements on this method.

#### Trainable Weka segmentation

Different image filters are used to generate image features as follows: Gaussian filer, Sobel filter, difference of Gaussians filter, Hessian matrix, Membrane projections and Median filter. TWS starts with semantic segmentation, which classifies each pixel into one of the three classes: background, foreground or edge. The class of foreground is of our interest. TWS uses *a random forest* classifier^38^ to determine a label for each pixel, i.e., a collection of decision trees where each tree represents a “test” if a given pixel contributes enough or not to the class (i.e., background, foreground or edge) prediction.

After we obtained all the foreground pixels, we added a step of instance segmentation to identify each single cell, such as giving each object an id. We used *connected component analysis*^*35,39*^ on the semantic segmentation to generate the indices.

#### Trainable Weka segmentation on GPU

Processing a single image, e.g., 4096 × 3000 pixels, takes ∼55 seconds when using TWS from Fiji^40^ (TWS provides a Fiji plugin for users without specific programming skills). This version runs on CPU. We use GPU to accelerate the whole process, and it takes ∼16 seconds (for segmenting an image of 4096 × 3000 pixels). We observed significant improvement in processing speed by implementing it on GPU. We implement GPUs (two NVIDIA QUADRO RTX 8000) to generate key image features by the RAPIDS cuCIM library^41^.

#### Reference dataset preparation (see Figure S4 for detail)

The reference image preparation was done with the Fiji image processing program^40^. We randomly selected 3 cropped images from the raw images and use the LabKit plugin^42^ (Weka-base segmentation) to generate the labeling of foreground, background and edge (edge labeling is optional) (Step 1-2, Figure S4). After that, the (foreground) segmentation result can be generated (Step 3, Figure S4) and converted to regions of interest (Step 4, Figure S4). The wrongly segmented objects can be manually corrected (Step 5, Figure S4) and the final segmentation result can be saved as reference ground truth data (Step 6, Figure S4). The whole process takes ∼30 min.

#### Random forest and its structure

Implementing TWS on GPU is one of our improvements to achieve fast and accurate high-throughput cell segmentation. The other improvement lies on the random forest optimization, which is introduced subsequently.

The nature of a random forest classifier indicates that how the forest ‘look alike’ and is critical for pixel-classifications. Variation (i.e., the parameters) includes the number of trees, the depth of each tree, and the number of nodes per tree, etc. If parameters of the model change, the prediction might also change, yielding the question “what are the best model parameters for our data of interest”?

To find the best random forest model per dataset, we applied grid search (and cross validation) to find the best parameters for the model, including the number of estimators, the maximum number of features, the maximum depth of the tree and the minimum number of sample leaf. To do this, a ground truth reference dataset is required, which, in our case, is a foreground/background labelled imaging dataset. This ground truth reference data acts as a reference during the process of selecting the best random forest parameter option (i.e., the depth of the tree and the number of sample leaf etc.) by checking the segmentation performance (i.e., accuracy or F1 score) (Figure 1, see Figure S3 for detail).

#### Key feature selection

We call the features that are relevant to our pixel classification the ‘key features’. Now the question comes to how to figure out the features that are the key ones. We applied *forward selection*^22^ to find out the key features - Recall that we have in total 57 default features *S* = {*I*_*j*_: *I*_*j*_ *is a feature image*, 1 ≤ *j* ≤ 57}, and we want to find a subset *s* ⊂ *S* as key features:

1. With respect to each *I*_*j*_ a classifier is trained, and corresponding segmentation accuracy is obtained via comparing to reference data.
2. The feature, say *I*_*k*_, that contributes to the highest accuracy will be selected as the first key feature *s* = {*I*_*k*_}. There are hence 56 features left (*S* = *S* − *s*), out of which we will choose the second key feature.
3. We let *s* = {*I*_*k*_, *I*_*i*_} where *I*_*i*_ ∈ *S* for every *i* ∈ {1, 2, …, *k* − 1, *k* + 1, …, 57}. A classifier is trained with every feature subset *s*, and the second key feature, say *I*,, is determined as *s* = {*I*_*k*_, *I*,} together contributing to the highest accuracy.

If we want to continue the selection, we shall repeat Step 2 by adding a third key feature *s* = {*I*_*k*_, *I*_*p*_, *I*_*q*_} where *I*_*q*_ ∈ *S* for every *q* ∈ {1, 2, …, *k* − 1, *k* + 1, *p* − 1, *p* + 1, …, 57}. The forward selection continues till the number of elements from *s* reaches a pre-defined number of features. For example, if the pre-defined number of features is 3, then the selection stops once we have *s* = {*I*_*k*_, *I*_*p*_, *I*_*q*_}.

In our case, using 3 key features provides us with the fastest and most accurate segmentation (Figure 2, S2 and S6).

### Training dataset preparation

For both MCF10A and GBM datasets, we selected 3 frames at different timepoints (frame 0, 360 and 720) from the entire time lapse movie (1080 frames in total, 4 min/frame). For annotation, we focused only at a partial region of a full field-of-view (FOV), and we manually annotated the chosen region with 3 classes, i.e., nuclei, background, and edge (Figure 1). All the annotation work was done with Fiji.

### Benchmarking

#### Input data

We randomly selected 5 cropped images from the MCF10A 3-day image data as our test images, and the ground truth on these test images were annotated manually. We compared our segmentation performance and computational time to other state-of-the art Weka-based cell segmentation methods. The methods that are compared to are listed as follows:

- *<u>TWS</u>*. This is the default setup of TWS, where there are 57 features generated for the random forest classifier (to determine if a pixel is of the Class foreground, background or edge).
- *<u>GTWS_whole features</u>*. Unlike TWS which uses a fixed structure of random forest (by default) for any input dataset, we search for the structure, out of many combinations, that can give the highest accuracy per input dataset. Optimization is bounded to the reference data, which helps to optimize the best parameters of the random forest classifier. In this case the whole features (57 features) are used. We aim to compare this method with TWS when the whole features are included during classification.
- *<u>GTWS_selected features</u>*. Instead of using the whole features, we apply only the key features (described above) to the classifier. Optimization of the random forest classifier, including key feature selection, is restricted to the reference data. We aim to compare this method to TWS when a subset of features is chosen.

The comparison results can be found in Figure 2.

To compare only the feature selection methods, we examine:

- *<u>Ilastik</u>*. Ilastik is a TWS-based cell segmentation method with an option of selecting key features. ilastik provides three kinds of feature selection methods^21^: filter method, GINI importance and wrapper method. According to the whole features *S* used for classifier, feature selection method helps users to choose a subset of it *s* ∈ *S*, on the classifier. We followed the filters which are provided by ilastik with sigma settings as 1, 2, 4, and 8. The used image filters are Gaussian filter, Laplace filter, gradient magnitude, difference of Gaussians, structure tensor eigen values and Hessian matrix eigen values. The total amount of image features provided in ilastik is 32. Please note that the image filters mentioned here are different than the filter setting used in the segmentation comparison analysis (Figure 2) as ilastik only offers the abovementioned filters (total 32 image features, compared to TWS or our GTWS methods, 57 image features). Also, for ilastik’s wrapper method, we set the parameter of size penalty to 0.01, 0.04, 0.1 and this method only allows for up to 6 features.
- *<u>GTWS_selected features</u>*. Here we use the same training dataset and image feature setting. We compare our GTWS methods with selected 3 key features with ilastik’s feature selection methods; our method outperforms ilastik’s feature selection methods.

The comparison results can be found in Figure S2, where we selected 3 to 9 features from the whole features.

### Segmentation performance

We examined the segmentation performance (F1 score) over different methods: TWS, ilastik and our GTWS method with whole features or with selected features. We visualize cell segmentation obtained from these methods in Figure 2. These outcomes were achieved using the same input image. Ground truth (GT) segmentation was prepared independently of all methods. We compared the segmentation outcome per method to the GT by calculating the IoU (intersection-over-union) value. We say a cell is well segmented if the calculated IoU ≥ 0.5, falsely segmented if 0.1≤ IoU < 0.5. When calculated IoU < 0.1 a cell segmentation becomes a false-negative. Here an IoU threshold is 0.5. The computational time was an average of six FOVs.

We quantify the segmentation performance (see Figure 2) by looking at the segmentation accuracy per method at the IoU threshold of 0.5. We obtain a TP when the calculated IoU ≥ IoU threshold, a FP if calculated 0.1 ≤ IoU < IoU threshold, a FN if calculated IoU < 0.1.

We observe that our GTWS method with whole features as well as with selected features have better performance than TWS and ilastik.

### Single cell tracking via FACT

The segmented cells are linked over time via the *Gaussian Mixture Mode*l-based method^2,43^.

#### Cell intensity as a Gaussian

First, one single cell and its intensity profile (Figure S9) is modelled as a 2D Gaussian, 𝒩(*x*; *μ, Σ*) with *x* be the 2D coordinates, *μ* be the mean location and Σ be the covariance (i.e., representing the shape of a Gaussian blob). Hereby we treat an object as a probabilistic model: Using the information of a cell at *t*, we could predict, for this cell, a “bounding box” at (*t* + 1), which in our case is a probabilistic distribution of (*u*^*t*^, *Σ*^*t*^). In other words, if we know the moving object distribution in the previous frame(s), we can locate the object in the next frame by tracking the expected changes. Tracking one cell means forwarding its Gaussian by calculating its expected changes, in the dimensions of location, shape and intensity.

#### Image as a Gaussian mixture

When dealing with an entire image, we want to forward all Gaussians from that image simultaneously over time. Then we move to the next step, modelling an image *I*^*t*^ (of many objects) at frame *t* as a Gaussian Mixture:

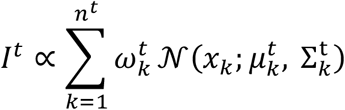

where *n*^*t*^ is the number of nuclei at frame *t, x*_*k*_ are the 2D coordinates for the *k*-th nucleus. The parameters 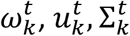, define the *k*-th Gaussian, i.e., respectively, they describe the contribution of the *k*-th nucleus to the image, estimated mean location and shape. Calculating the expectation per cell over time is performed by a full Bayesian approach^2,24^, with the assumption that cell properties between two consecutive frames are correlated.

#### Cell division

We handled cell divisions separately in this process: First we shall determine if a cell is going to divide, then we include the newly generated cell into a Gaussian mixture. In Figure S10 we give an illustration of determining a dividing event: For each nucleus we perform Otsu thresholding on only the foreground pixels, and we see if two unconnected regions are generated. If not, we do not find a case of dividing and we do not proceed with splitting one Gaussian into two components. If two unconnected regions are found, we further assess if the size per region is larger than a pre-defined value (as a region can be too small to represent a daughter cell).

Once we have detected a cell division at frame *t*, we accordingly need to update the Gaussian mixture at *t*, by letting *n*^*t*^ = (*n*^*t*^ + 1).

### Cell track correction

#### Solution to merging-demerging problem

We propose to correct the corrupted tracks by three steps (see Figure 5a):

- Step 1: We examine if a cell division is valid. A cell division is a false-positive if it is next to one or more incomplete tracks in both space and time. In Figure 5a, a merging at *t* = 2 causes a cell track (orange) disappearing at location *x*_*disappear*_, a demerging at *t* = 3 causes a division at location *x*_*divide*_. We call *x*_*disappear*_ also the location of the incomplete track (orange). The cell division is next to the incomplete track in space if the (Euclidian) distance Δ*x* between *x*_*disappear*_ and *x*_*divide*_ is smaller than a pre-defined value (such as 10 pixels). The cell division is next to the incomplete track in time if the time difference Δ*t* between the merging and demerging events is smaller than a pre-defined value (such as 5 frames). In this example, the cell division is found invalid, and we move to the next step.
- Step 2: For an identified false division, we cut the link between the mother cell and one of the two daughter cells. The daughter cell that is closer to the incomplete track is excluded. We remove the link (green) from *t* = 2 to *t* = 3, as this daughter (green) is closer to *x*_*disappear*_ than the other daughter (blue) is. The link removal generates an incomplete track (green) from *t* = 3 to *t* = 4.
- Step 3: We connect two incomplete tracks, and close the gap between the two incomplete tracks. In Figure 5a, the two incomplete tracks (orange and green) are to be connected. There is a gap between them, which refers to the disappeared cell at *t* = 2. We then construct a fake cell (gray) at this frame at a random location *x*_*generate*_ between *x*_*disappear*_ and *x*_*divide*_. The time difference Δ*t* = 1 indicates one fake cell is required. We update the reconstructed cell to the subsequent frames *t* = 3, 4. At *t* = 4 we obtain two complete tracks.

### Tracking accuracy and computational time

#### Tracking accuracy

We assess the accuracy of cell linking by examining how many cell tracks, out of all, are accurately followed. We call a cell is accurately tracked if its centroids between any two adjacent time points are accurately linked: Let (*x*_*i*_, *y*_*i*_, *t*) represent the centroid coordinates of cell *i* at *t* ∈ {1,2, …, *T*} with *T* be the length of a time lapse movie. Cell track *i* is accurate if (*x*_*i*_, *y*_*i*_, *t*) is accurately linked to (*x*_*i*_, *y*_*i*_, *t*+1) for every *t*.

In practice we chose two small regions for manual checking the detected cell tracks:

- Region 1 (of dimension 1170 × 724 pixels): There are 354 cell tracks, out of which, 17 tracks are contaminated with at least one frame being wrongly linked. This gives precision of 95%.
- Region 2 (of dimension 1628 × 978 pixels): There are 356 cell tracks, out of which, 19 tracks are contaminated with at least one frame being wrongly linked. This gives precision of 94%.

The tracked images are shown in Supplementary Information: Supplementary Video 10 for Region 1 and Supplementary Video 11 for Region 2.

#### Tracking efficiency

For real-time experiment, the tracking efficiency is crucial. A real-time tracking method shall be able to catch up the speed of image generation. In our experiment, tracking two frames of ∼20,000 cells per frame took ∼100 seconds (i.e., an average of three tests outcomes, 101.37, 98.50, and 99.96 seconds).

### Cell lineage accuracy

We calculate, for each lineage tree, its F1 score: we check manually if each detected division from this tree is a TP or FP. We also check any missed division (compared to the raw imaging) and treat it as a FN. We define that a detected division is a TP if 1) it is truly a division from this lineage, as well as 2) the detected splitting time is +/-5 frames compared to the true time. For example, if a detected lineage tree has 5 divisions, with 2 divisions being FPs, 3 divisions being TPs, and 1 missed division as 1 FN, then the F1 score of this detected tree is 0.67 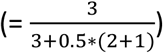.

In Table S1 the performance of lineage tracking is shown. On average we can reach 91% F1 score per lineage.

## Acknowledgements

MPC acknowledges support from the Oncode Institute, Cancer GenomiCs.nl (CGC), NWO (the Netherlands Organization for Scientific Research) Veni and Vidi Grant, Stichting Ammodo and Erasmus MC grant. MPC appreciates Josephine Nefkens Stichting’s support on the UFO microscope.

## Author contributions

TCC scripted the cell segmentation algorithm and analyzed the assayed data. LY scripted the real-time cell tracking algorithm and analyzed the assayed data. CB contributed to the cell culture preparation for the experiments. KF contributed to the GBM cell preparation. TCC, LY and MPC designed all the experiments. MPC, LY and TCC wrote the paper with input from all authors. MPC initiated the project and contributed to and supervised all aspects of the project.

