## Supplementary Information for "Fast and Accurate Cell Tracking: a real-time cell segmentation and tracking algorithm to instantly export quantifiable cellular characteristics from large scale image data"

Supplementary Figure S1-S10
Supplementary Table S1
Supplementary Methods
Supplementary Video 1-11

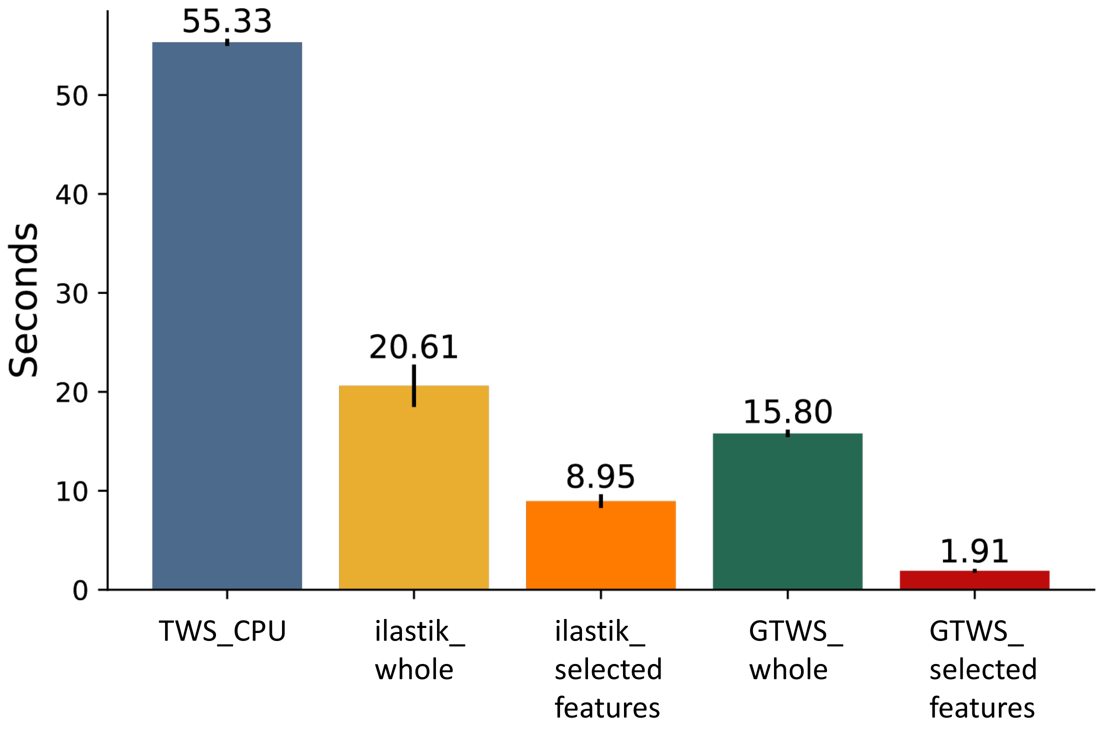


**Figure S1. Computational time comparison.** Computational time for TWS segmentation using CPU and GPU (only our GTWS methods use GPU). Please note that ilastik uses a total of 32 image features (ilastik_whole) and TWS_CPU and our GTWS_whole methods use a total of 57 image features. “Selected features” indicates the computation uses the selected 3 key features.

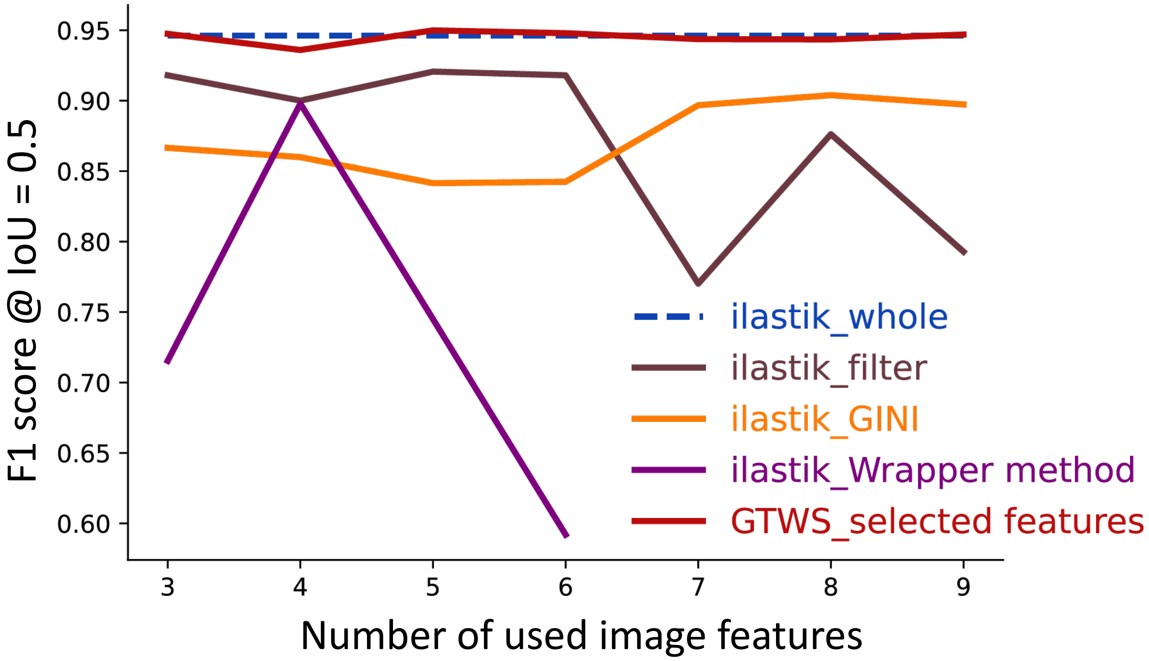


**Figure S2. Features selection comparison.** Segmentation performance (F1 score) for different feature selection methods with various amounts of used features is shown. The compared methods include our own GTWS' top feature selection method (GTWS_selected features), ilastik's methods using the whole default 32 image features (ilastik_whole) or ilastik's feature selection methods (ilastik_filter, GINI, Wrapper method). Please note that the image features we use here are from ilastik's default setting (32 image features), which are different than the whole image features used in TWS and our GTWS methods (57 image features, see Materials and Methods for detail), therefore the data of using TWS' and GTWS' whole features are not shown here. Please also note that ilastik's wrapper method only allows for the use of up to 6 image features.

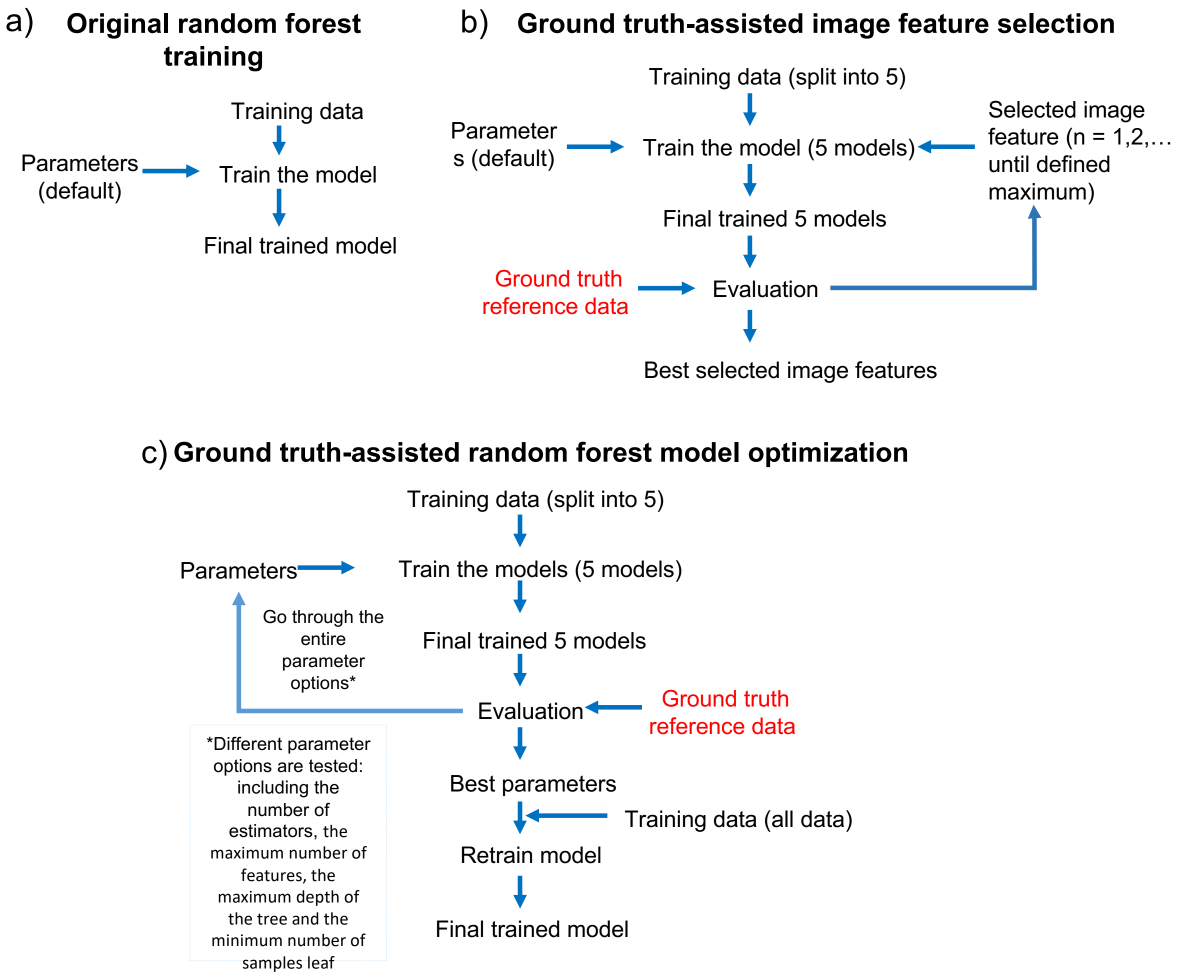


**Figure S3**. **Ground truth-assisted model training.** (a) Original random forest training: all the training data will be trained once using the default parameter setting. (b) Ground truth-assisted image feature selection. Training data are split into 5 and trained separately using the default parameter setting. The best feature combination will be selected with the forward selection method. Users can define the maximum amount of image features to be used for classification. In our GTWS method, 3 features are the default setting. The feature selection procedure starts with identifying the best one feature, evaluated by the ground truth reference data (based on segmentation performance). The following 2$^{nd}$ and 3$^{rd}$ best feature (or whatever the maximum feature number defined by users) are also selected sequentially by evaluating with the ground truth refence data (based on segmentation performance). (c) Ground truth-assisted random forest optimization. Training data (either whole image features or selected features) are split into 5 and trained separately using one of the parameter options (162 options when using whole features, 27 options when using selected features): including the number of estimators, the maximum number of features, the maximum depth of the tree and the minimum number of samples leaf). The final trained 5 models will be evaluated using the ground truth reference data and repeat the procedure until all the parameter options have been tested. The best parameters for the random forest classification model will be selected by comparing the result through evaluating with the ground truth reference data (based on segmentation performance). The best parameters will then be used to retrain the random forest model to obtain the final, best one.

Step 1: Load the raw reference image with Fiji. Open Labkit. (Plugins >> Labkit >> Open current image with LabKit )

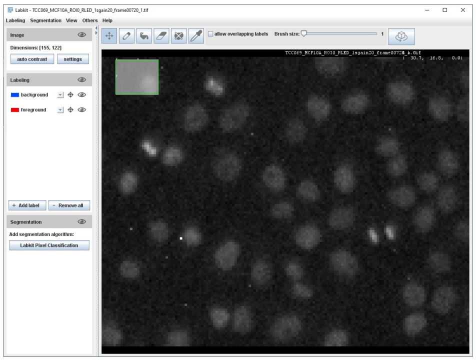

Step 2: Add the training data for each labeling (foreground, background, edge). Then, click the Labkit Pixel classification.

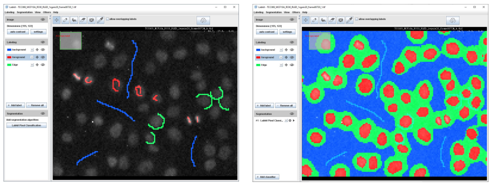

Step 3: Get the segmentation result from Labkit (Segmentation >> Create label from segmentation >> foreground )

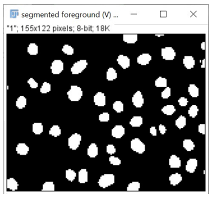


(Optional step) Add one more label, edge, during labeling

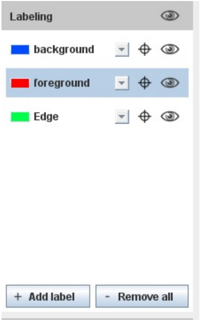


Step 4: Convert the segmentation result to regions of interest (ROIs) (Plugins >> BIOP >> Image Analysis >> ROIs >> Label image to ROIs)

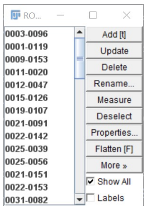


Step 5: Manual correction on ROIs. For example, the ROI15 of the following image (shown in blue) are connected to each other and can be corrected manually (by re-drawing the cell counter by users). The corrected image is shown in the following right figure.

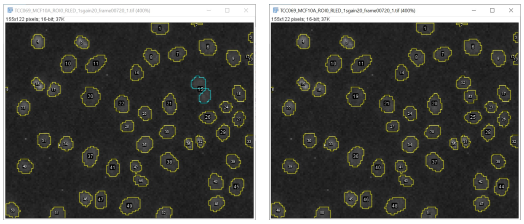

Step 6: Generate the segmentation result and save it to a 16-bit image (Plugins >> LOCI >> ROI Map)

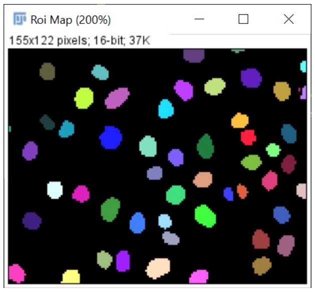


**Figure S4. Generation of reference ground truth data**. 6 steps (Step 1-6) to generate reference ground truth data via Fiji are described.

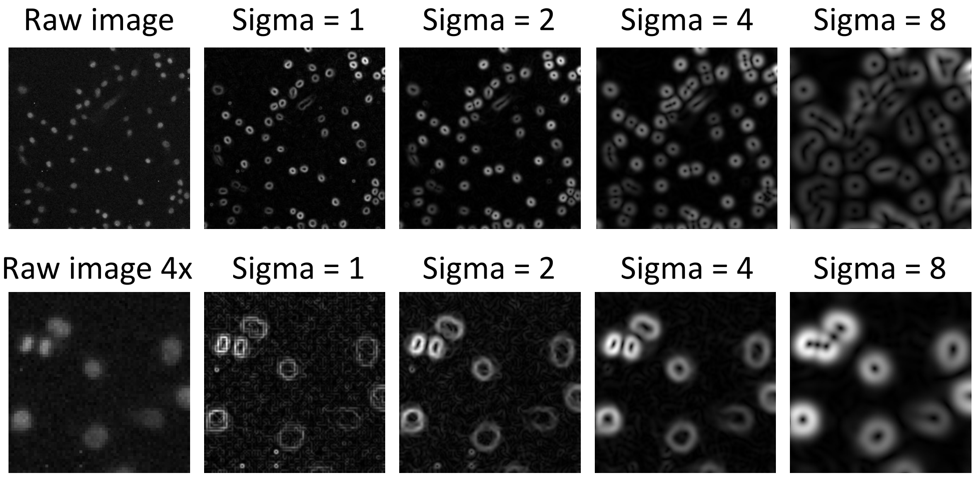


**Figure S5**. **Image features. Top panel**, from left to right: a raw image of MCF10A cells (nuclear size is ~ 15x15 pixels) and its processing results with Sobel filtering using a sigma value of 1, 2, 4 or 8. As shown, processing using Sobel filtering with a sigma value of 1 and 2 enhanced the edge of the nuclei accurately, but not with a sigma value of 4 and 8. **Bottom panel**, from left to right: a zoomed-in raw image from the top panel (4x zoom, nuclear size is ~ 60x60 pixels) and its processing results with Sobel filtering using a sigma value of 1, 2, 4 or 8. As shown, processing using Sobel filtering with a sigma value of 2 and 4 enhanced the edge of the nuclei better than using a sigma value of 1 and 8. This indicates that not all the image features are relevant for classification.

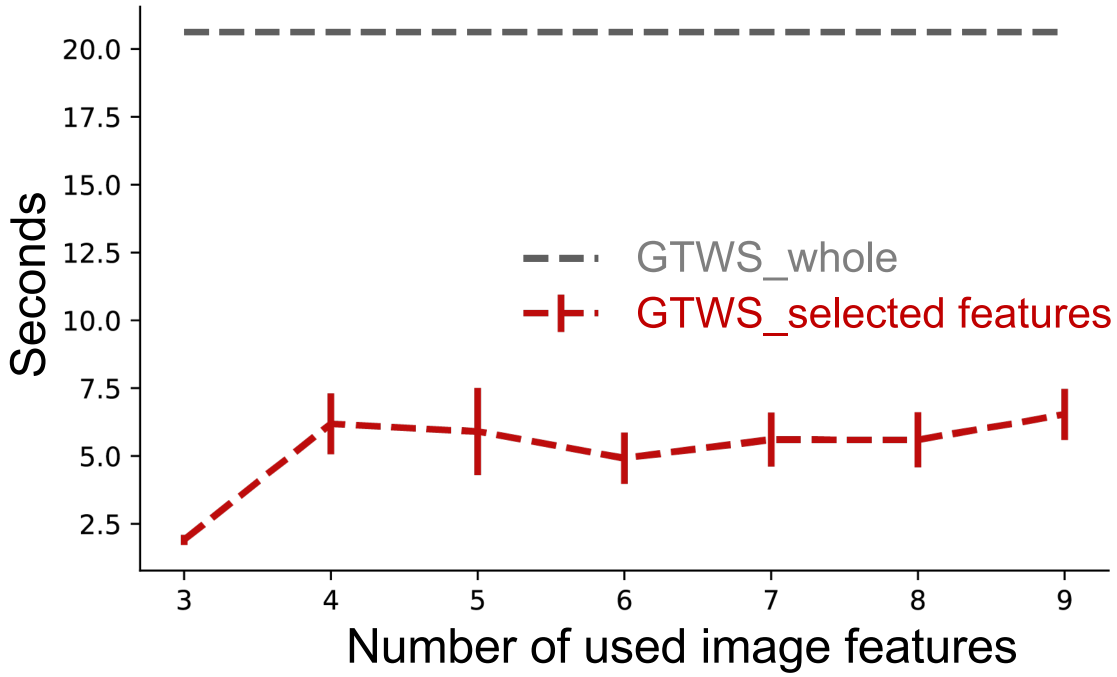


**Figure S6. Computational time with different numbers of used image features.** Computational time for our GTWS segmentation using whole features (gray line) or selected features (red line) (3 to 9 key selected features were used).

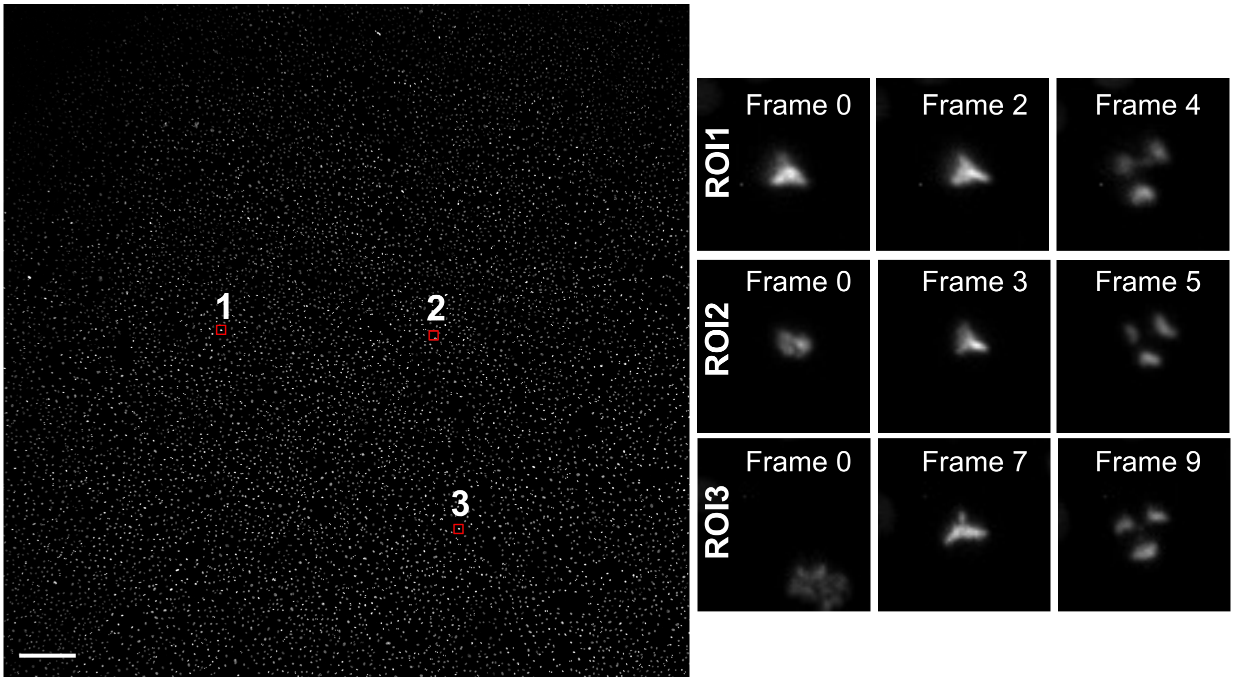


**Figure S7. One representative field-of-view (FOV) of tripolar division detection.** Left panel: A full view of a 5120 x 5120 pixels image with ~15,000 MCF10A breast cancer cells (stained with a SPY650-DNA nuclear dye). The recorded frame rate is 10 min/frame. The red boxes (region-of-interest (ROI) 1-3) show the locations of the occurring tripolar dividing cells during the recording (within 1.5 hours). Right panel: Zoomed-in images of the highlighted ROIs where tripolar division occurred. The Frame number indicates the timing of the occurrence of tripolar division. Scale bar: 500 μm.

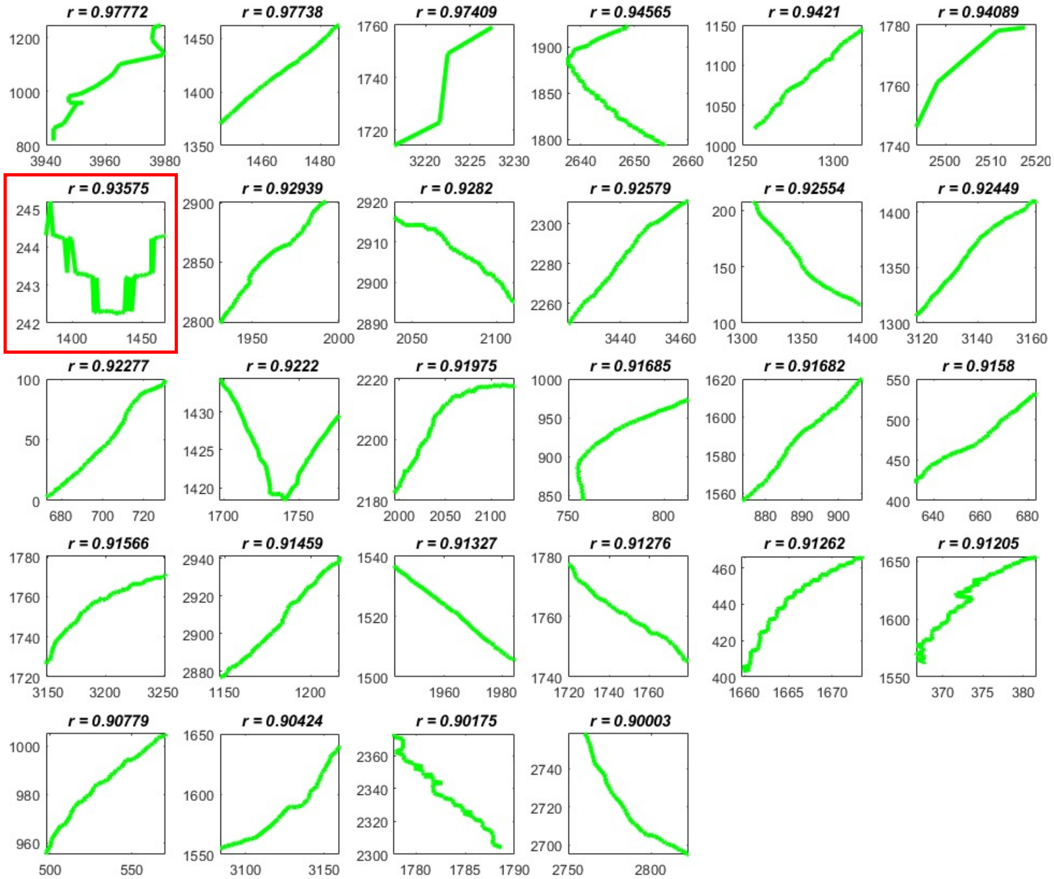


**Figure S8. 28 cell trajectories with directionality ratio > 0.90.** We observed 1/28 were falsely detected (highlighted in red) as directional walk.

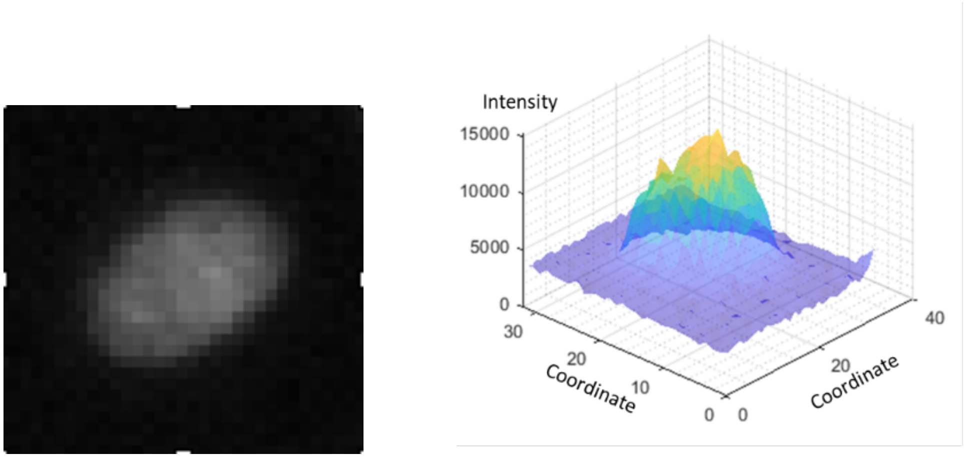


**Figure S9**. **2D Gaussian modeling of a cell.** Raw image of a single cell (left) and its intensity profile (right). The intensity can be modelled as a 2D Gaussian.

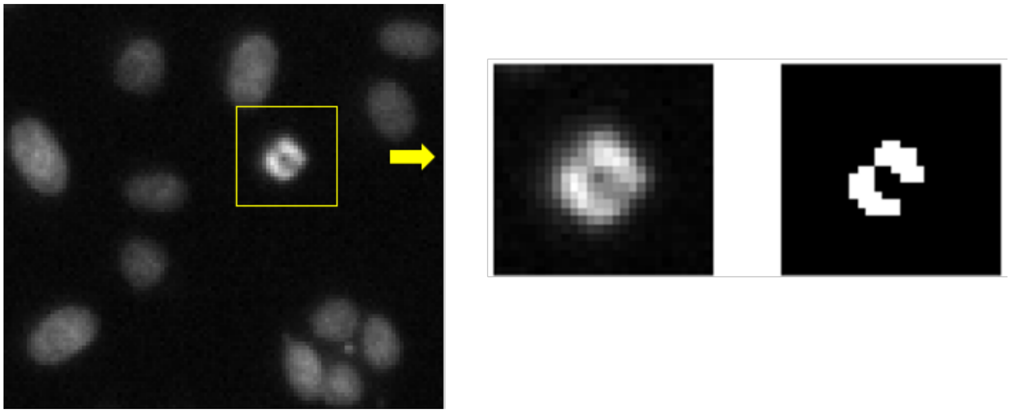

**Figure S10**. **Dividing event and detection.** To see if a cell (highlighted, left) is about to divide, we apply thresholding on the foreground pixels (middle) and assess the number of unconnected regions (right). In this case two unconnected regions are generated, hence this Gaussian is splitable.

**Table S1.** Detected lineages with accuracy check-up. For each group of detected trees, we randomly picked a certain number of trees for assessing accuracy. We take into account the cases of TPs, FPs and FNs. A detected division is a TP if 1) it is truly a division from this lineage, as well as 2) the detected splitting time is +/- 5 frames, compared to the true division time. We show F1 ratio (= $\frac{\#TPs}{\#TPs + 0.5*(\#FPs + \#FNs)}$).

| **Trees** | **Tree-1-div** | **Tree-2-div** | **Tree-3-div** | **Tree-4-div** | **Tree-5-div** |
|---|---|---|---|---|---|
| # Trees (detected) | 2705 | 625 | 353 | 27 | 4 |
| # Trees (checked) | 27 (1%) | 13 (2%) | 17 (5%) | 13 (50%) | 4 (100%) |
| F1 ratio | 0.98 | 0.92 | 0.97 | 0.87 | 0.83 |

### Supplementary Methods

#### Cell Culture

MCF10A-H2B-GFP cells

MCF10A-H2B-GFP breast epithelial cells, a gift from Dr. Reuven Agami (Dutch National Cancer Institute, NKI), were grown in DMEM/F-12 (ThermoFisher) supplemented with 5% horse serum, 1% penicillin/streptomycin, Epidermal growth factor (10 ng $ml^{-1}$; ThermoFisher), Hydrocortisone (500 ng $ml^{-1}$; Stem Cell), cholera toxin (100 ng $ml^{-1}$; Sigma) and insulin (10 µg $ml^{-1}$; ThermoFisher) in a 37 °C incubator under 5% CO2.

Before conducting experiments, ~50,000 cells were seeded on a fibronectin (0.1 mg/mL)-coated 35 mm-glass bottom dish with a 20 mm-microwell (Cellvis) in the MCF10A culture medium (described above) without phenol red and were stained with a SPY650-DNA nuclear dye (Spirochrome) (1:2000) ($\lambda_{ex}$: 652, $\lambda_{em}$: 674nm). Experiments were performed ~24 hours after plating on the glass-bottom dishes.

Glioblastoma (GBM) cells

GBM cells were maintained on reduced growth factor basement membrane extract (Cultrex, Bio-techne R&D systems) coated petri dishes in serum-free Dulbecco's Modified Eagle Medium (DMEM)/F12 supplemented with 1% penicillin/streptomycin, 2% B27 without vitamin A (Gibco, ThermoFisher), 20ng/mL basic fibroblast growth factor (Gibco, ThermoFisher), 20ng/mL epidermal growth factor (Sigma Aldrich), and 5µg/mL heparin (Alfa Aesar, ThermoFisher). Cells were maintained in a humidified incubator (37°C, 5% CO2) and culture medium was refreshed every 3-4 days). Passaging of the cells was performed around 80-90% confluency using enzymatic dissociation (Accutase, Invitrogen). The use of GBM samples and study was approved by Erasmus University Medical Center ethics committee (MEC-2013-090).

Before conducting experiments, ~19,000 cells were seeded on Cultrex (0.1 mg/mL)-coated 35 mm-glass bottom dish with a 20 mm-microwell (Cellvis) in the GBM culture medium (described above) without phenol red and were stained with a SPY650-DNA nuclear dye (Spirochrome) (1:2000) ($\lambda_{ex}$: 652, $\lambda_{em}$: 674nm). Experiments were performed ~24 hours after plating on the glass-bottom dishes.

### Supplementary Videos

**Supplementary Video 1.** Time lapse movie from Main text Figure 4b, top panel.

**Supplementary Video 2.** Time lapse movie from Main text Figure 4b, bottom panel.

**Supplementary Video 3.** Time lapse movie from Main text Figure 5b.

**Supplementary Video 4.** Time lapse movie from Main text Figure 5c.

**Supplementary Video 5.** 5 hr-time lapse movie of GBM cells.

**Supplementary Video 6.** 24 hr-time lapse movie of MCF10A cells.

**Supplementary Video 7.** 'tree-3-div' cell lineage time lapse movie.

**Supplementary Video 8.** 'tree-4-div' cell lineage time lapse movie.

**Supplementary Video 9.** 'tree-5-div' cell lineage time lapse movie. Main text Figure 7b.

**Supplementary Video 10.** Time lapse movie used for checking tracking accuracy (Materials and Methods, Main text).

**Supplementary Video 11.** Time lapse movie used for checking tracking accuracy (Materials and Methods, Main text).
